# Intensity Dependent Inhibition of Single Pulse TMS on Stretch-Evoked Long-Latency Responses in the Flexor Carpi Radialis

**DOI:** 10.1101/2024.03.18.585555

**Authors:** Cody A. Helm, Fabrizio Sergi

## Abstract

Transcranial magnetic stimulation (TMS) can modulate corticospinal excitability during stretch-evoked long-latency responses (LLR). It has been previously established that *suprathreshold* TMS intensities can partially inhibit the cortical contribution to LLRs relying on a cortical silent period occurring after TMS. However, it is unknown whether the TMS-induced inhibition of stretch-evoked LLRs that relies on the cortical silent period can also be achieved via *subthreshold* stimulation.

In this study, twelve healthy participants performed a protocol combining surface electromyography (EMG), robot-evoked wrist perturbations, and single pulse TMS applied to the motor cortex to study the effect of TMS intensity on the LLR amplitude in the flexor carpi radialis. We tested two TMS intensities of *subthreshold* (90%) and *suprathreshold* (130%) of the active motor threshold applied such that the motor evoked potential (MEP) peak would arrive 50 ms prior to perturbation onset.

In *suprathreshold* TMS trials, TMS significantly reduced the cortical contribution to a LLR when applied prior to perturbation onset. When comparing the effects measured in the presence and absence of robot-applied muscle stretch, we observed that only *suprathreshold* conditions achieved inhibition of LLR, while *subthreshold* conditions did not result in any inhibition. Overall, our findings establish a clear distinction between the effect of *subthreshold* and *suprathreshold* TMS on the LLR inhibition via the cortical component of the silent period.

## INTRODUCTION

The human neuromuscular system incorporates feedback mechanisms to continuously update movements as we receive new sensory information from our environment. One feedback mechanism is the stretch response which is defined as a burst of EMG activity following an imposed limb displacement (1, 2). An important component of the stretch response is the long-latency response (LLR), which occurs from 50 ms to 100 ms following perturbation onset (3). The LLR occurs between the purely spinal short latency response and the delayed voluntary reaction, each occurring 20 ms to 50 ms and more than 100 ms following perturbation onset, respectively. Notably, the LLR has both spinal and supraspinal components (4, 5, 6).

Multiple motor pathways contribute to the generation of the LLR. A large body of evidence shows that the primary motor pathway, the corticospinal tract (CST), contributes to the LLR (7, 8, 4, 9, 6, 3, 10, 11, 1, 12, 13, 14, 15, 16). Strong evidence for the involvement of the CST in the LLR stems from the association between motor cortex neural activity and the long-latency response as well as transcranial magnetic stimulation paradigms (13, 10). Additionally, the reticulospinal tract (RST), a secondary motor pathway, has been suggested to contribute to the long-latency response (17, 18, 19, 20). Research with nonhuman primates has shown the RST to be associated with hand grasping following corticospinal and rubrospinal lesions (21, 22, 23). Excitatory RST connections have been found to hand motoneurons in nonhuman primates and may be as strong as RST connections found in the more proximal muscles (24). In humans, current evidence suggests that the RST is responsible for the increase in the long-latency response when participants are instructed to resist a perturbation rather than yield, highlighting that it contributes to the task-dependent changes (3, 18).

Using functional MRI, neural correlates of the LLRs were observed in the brainstem during wrist perturbations with the StretchfMRI paradigm (25). These studies provide the muscle-specific brainstem areas associated with the long-latency response and support the notion that the RST contributes to the long-latency response. Additionally, the StartReact experimental paradigm has shown to elicit a faster than normal planned voluntary muscle activity (∼ 70 ms post perturbation onset) in the presence of a startling acoustic stimulus prior to a perturbation (26, 27). The decrease in the voluntary response latency is likely contributed by the RST as the only supraspinal pathway able to act with such a short response time (28, 26, 18). Furthermore, a joint perturbation and a startling acoustic stimuli induce a similar EMG recorded response, suggesting that both responses may be generated by the same subcortical pathway (29).

While our understanding of the motor function of subcortical pathways like the RST is improving, due to limitations in the methods used in previous studies, it is still currently unknown if the RST can generate stretch reflex activity independently from the CST, or if the RST acts solely as a relay for descending CST signal. In the former case, the RST could generate functional motor responses, stretch reflexes, also in absence of CST input. This would support the hypothesis that the RST can provide functional motor responses, regardless of CST input, suggesting its beneficial role in motor recovery in the presence of CST damage. Therefore, alternative methods need to be developed to study the isolated role of the RST on LLR formation, whereby a causal role of this pathway in producing LLR can be identified.

One way for studying the role of subcortical pathways on LLR is to identify their contribution under different levels of involvement of the primary motor pathway, for example by modulating cortical excitability. Transcranial magnetic stimulation (TMS) is a neuromodulation technique that can be used for this purpose. When TMS is placed over the motor cortex, motor evoked potentials (MEPs) are measured via surface electromyography of relevant muscles, with a time delay comprised between 10 ms and 30 ms. Following the MEP, there is a decrease in the muscle activity of the targeted muscle which has been referred to as the silent period (30, 31, 32, 33, 34, 35). This TMS-induced silent period lasts approximately 150 ms to 200 ms, with the first 80 ms of inhibition coming from spinal inhibition and the later part coming from longer lasting cortical inhibition via GABA receptors (35, 36, 37, 38). As such, TMS can modulate cortical excitability, and thus modulate the contribution of cortical pathways during a stretch-evoked motor response, which would allow to study the contribution of secondary motor pathways to LLRs during reduced cortical excitability. With appropriately timed TMS pulses, we can overlay the cortical component of the silent period with the LLR to specifically target the cortical portion of the LLR as oppose to spinal inhibition.

The capability of TMS to inhibit the stretch-evoked long-latency response has been studied under a variety of TMS intensities, latencies between evoked MEP and perturbation onset, and for a variety of tasks. The previous studies that use TMS in the attempt to modulate LLRs are displayed in Fig. 1, broken down by the parameters of TMS stimulation (TMS amplitude and timing), and with an indication of what the measured effects have been.

**Figure 1.**
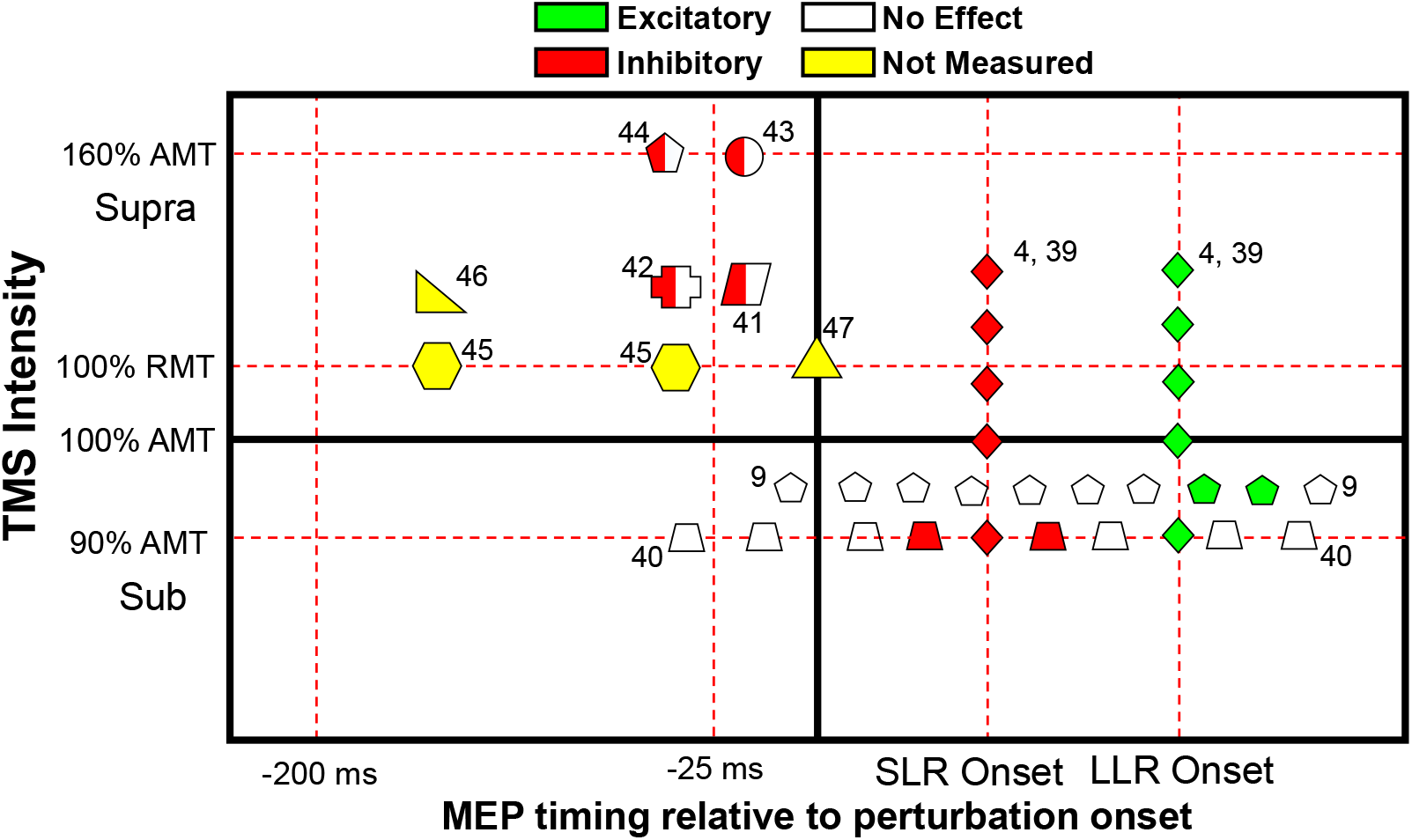
Schematic of previous LLR studies involving TMS. TMS intensity relative to motor threshold is labeled on the y-axis. The timing of the ensuing motor-evoked potential (MEP) relative to perturbation onset is labeled on the x-axis. Each shape corresponds to a different study. Green indicates that the effect of TMS on the LLR was excitatory. Red indicates that the effect of TMS on the LLR was inhibitory. White indicates there was no effect of TMS on the LLR. Yellow indicates the study did not measure the effect of TMS on the LLR.

When timed to evoke an MEP just post perturbation onset, *suprathreshold* TMS (100% - 140% AMT) reduces the LLR amplitude when the MEP peak arrives during the short-latency response onset (39, 4). When *suprathreshold* TMS is timed such that the evoked MEP arrives during long-latency response onset, there is an increase in the EMG response that is greater than the sum of the MEP and LLR alone (39, 4). Additionally, when *subthreshold* TMS (96% AMT) is timed such that the ensued motor response arrives 60 ms to 90 ms post perturbation onset, there is an increase in the LLR when the TMS-induced short term facilitatory EMG is overlayed with the long-latency response (9). This demonstrates an increase in the cortical excitability when the MEP is timed to coincide with the LLR window. Additionally, when *subthreshold* TMS (90% AMT) is applied such that no short-term facilitatory EMG is observed, there is a reduction in the LLR amplitude when applied between 55 ms and 85 ms prior to LLR onset in the tibialis anterior muscle (i.e., MEP peak would occur 35 ms to 65 ms prior) (40). However, these studies likely achieve inhibition primarily via the *spinal* component of the silent period.

When timed to evoke an MEP occurring prior to perturbation onset, *suprathreshold* TMS (120% resting motor threshold) has been shown to suppress the LLR anticipatory modulation associated with a multi-joint dynamic mechanical environment, but not the LLR itself (41). Furthermore, at the same timing, *suprathreshold* TMS inhibits the LLR increase associated with a resist task instruction in the flexor carpi radialis (42). However, previous work also found that a strong *suprathreshold* TMS, (160% AMT), arriving prior to perturbation onset, reduces the LLR amplitude but not the LLR modulation associated with the environmental stiffness (43, 44). The remaining TMS studies in Figure 1 apply TMS to the motor cortex but do not quantify the excitatory or inhibitory effect of TMS on the LLR amplitude (45, 46, 47).

For studies that aim to use TMS to decouple the contribution of cortical and subcortical circuits to LLRs, and aim to use low temporal resolution imaging methods to measure the contribution of subcortical circuits in presence of cortical inhibition, it would be ideal to use the smallest possible TMS amplitude, and to time the TMS stimulus such that the possible MEP would arise before perturbation onset. If *subthreshold* TMS can inhibit the LLR in a similar manner to *suprathreshold* TMS, its application would be beneficial as it reduces the excitatory/confounding effect of TMS stimulation on the motor cortex regions, as would be observed with *suprathreshold* TMS. Additionally, TMS timed such that the MEP would arrive prior to perturbation onset would achieve the expected inhibition by primarily engaging the cortical, and not spinal, component of the TMS-evoked silent period during the LLR, allowing a clean interpretation of the imaging outcomes.

Unfortunately, previous studies achieving LLR inhibition via TMS by the cortical silent period mechanism did not pursue an experimental design that allowed to compare the effects across multiple stimulation amplitudes. As such, we don’t know if *subthreshold* TMS can inhibit the cortical contribution to LLRs.

To our knowledge, no studies have investigated the intensity-dependent effect of TMS-induced inhibition on LLR responses, when the TMS was timed to modulate primarily the cortical component of the silent period. The objective of this study was to determine the effect of TMS intensity on its capability to reduce the CST’s contribution to the LLR in a forearm flexor muscle, FCR. Additionally, we wanted to evaluate the accuracy of TMS coil placement and stimulation of the forearm motor hotspot with stimulation marker tracking. The outcomes of this study provide insight into the mechanisms by which *suprathreshold* and *subthreshold* TMS result in a reduction of LLR via the cortical component of the EMG silent period in the forearm. The knowledge from this study may be used to study the effect of cortical inhibition on the LLR, and may inform the design of neuroimaging studies that aim to quantify brain function associated with LLRs.

## MATERIALS AND METHODS

### Participants

Twelve healthy participants (6 F, age range: 19–33 years) performed the experimental protocol. The participants were exposed to the same TMS modes. All of the participants were right-handed and free from neurological disorders and did not have any contra-indications to MRI or TMS. The subjects provided informed written consent for the study. The study was approved by the Institutional Review Board of the University of Delaware (IRB no: 1545543-8, and -9) and was in accordance with the Declaration of Helsinki.

### Experimental Setup

In this study, we combined a wrist robot, the MR-StretchWrist, used to elicit stretch reflexes in the forearm, with TMS, to stimulate the motor cortex with precise timing, and surface EMG, to record muscle activity in one forearm muscle. Details on these elements are provided below.

### Surface Electromyography

Bi-polar surface electrodes were placed on the skin over the flexor carpi radialis (FCR) to record muscle activity. The location of the FCR muscle belly was determined by having the subject flex and relax their wrist and palpating down the forearm muscle. The skin was cleaned with a 70% Isopropyl alcohol wipe (Med Price) to remove any dirt and debris on the surface of the forearm. A thin layer of skin was abraded with Nuprep skin prep gel (Weaver and Company) to reduce contact impedance. The electrodes (standard multitrode Ag/AgCl bipolar electrodes from Brain Products, Munich, Germany) were firmly attached on the skin surface using double-sided adhesive collars (BIOPAC Systems, Inc.). The bi-polar electrodes were aligned with the direction of the muscle fibers. An additional electrode (ground electrode) was placed on the olecranon in the back of the elbow to serve as a ground. Each electrode was filled with a high-chloride abrasive electrolyte gel (EASYCAP) to further reduce contact impedance. Muscle activity was amplified using an 8-channel bipolar amplifier (MR Plus ExG, Brain Products, Munich, Germany). Muscle signal and external triggers were synced together using a Trigger and Sync Box (Brain Products). The EMG signal was visualized and recorded using BrainVision Recorder software. Proper electrode placement was confirmed by having the subject perform flexion and radial deviation contractions and observing a high signal to noise ratio in agonist muscles.

### Robotic perturbations

We used the MR-StretchWrist to apply wrist perturbations and evoke stretch reflexes on forearm muscles. The MR-StretchWrist is an MRI-compatible wrist robot previously developed by the Human Robotics Lab (25) at the University of Delaware. The robot uses a piezoelectric ultrasonic motor to apply velocity-controlled perturbations to the wrist joint and an incremental encoder to continuously measure wrist angle. The MR-StretchWrist incorporates a force/torque sensor to measure wrist torques applied by the subject.

During a wrist perturbation, a velocity command of 150^*°*^/sec is sent to the MR-StretchWrist for 200 ms in the extension direction to elicit a stretch reflex response in wrist flexors, including the FCR muscle. Following perturbations, the wrist robot returns the wrist to the neutral posture at a slow velocity (35^*°*^/s) to avoid eliciting a stretch reflex.

### Transcranial Magnetic Stimulation

A neuronavigation procedure was performed to position the TMS coil over the participant’s forearm motor hotspot of the motor cortex. First, the participant’s structural brain scan was loaded into the Neuronavigation software (Localite 3.3.0). With the Localite software, brain segmentation was performed to virtually extract a 3d rendering of the brain volume from the skull. We converted from subject space to Talairach space by manually defining the anterior commissure, posterior commissure, and cortical falx point on the brain scan. A target marker was placed on the forearm motor hotspot of the motor cortex as the location that we aimed to stimulate with TMS, which was confirmed to be around (-20, -20, 50) in MNI space. The target marker was defined visually by identifying the pre-central gyrus, and moving laterally along the gyrus until reaching the hotspot for muscles responsible for movements of the distal upper extremity. An entry marker was placed on the surface of the skull where the TMS coil would be positioned. The entry marker was oriented with the TMS coil handle pointed posteriorly, approximately 45^*°*^ from the mid-sagittal line for adequate target stimulation to occur (48). See Fig. 2 for the location of the target, entry, and TMS instrument coil from an example participant.

**Figure 2.**
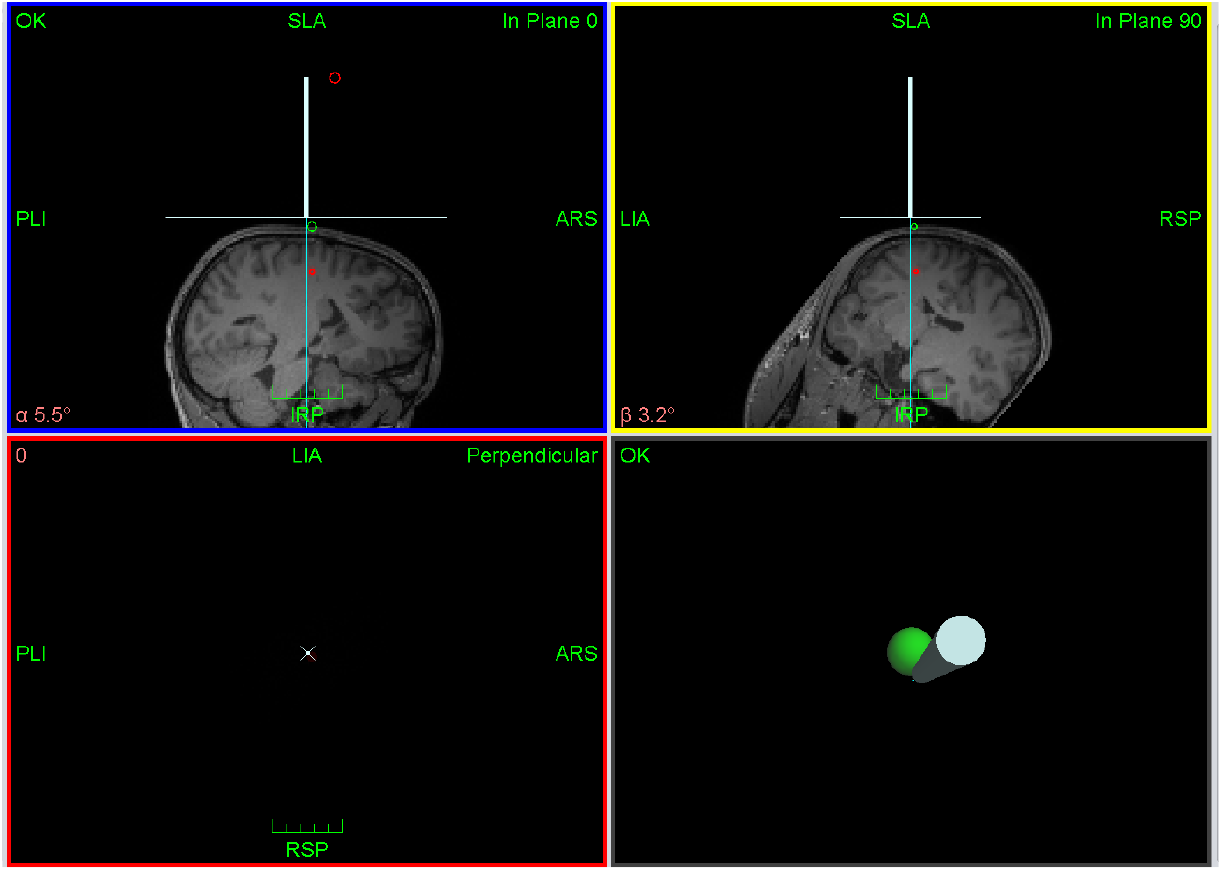
Localite Neuronavigation software displaying the position and orientation of the TMS coil (blue line) and the position of the entry marker (green circle) and target marker (red circle) for accurate coil placement. Blue box displays the sagittal plane and the yellow box displays the coronal plane. The bottom right box displays the plane looking down on the entry-target axis and the red box displays the plane perpendicular to the entry-target axis plane.

Next, we performed participant registration to register the physical features of the participant’s head with the Neuronavigation system. We used motion capture tracking and reflective markers to register the position and orientation of the participant’s head and TMS coil in 3D space. A figure-of-eight TMS coil (MCF-B65, Magventure) was calibrated to the camera which defined the position and orientation of the center of the TMS coil relative to the coil’s reflective markers. We placed reflective markers on the participant’s forehead via double-sided skin adhesive tape for participant registration. Participant registered included a ‘landmark registration’ and a ‘surface registration’. For ‘landmark registration’, the ‘left tragus’, ‘right tragus’, and ‘nasion point’ were manually defined on the brain scan. A pointer was used to calibrate the features of the participant’s head with the respective markers on the brain scan. During ‘surface registration’, a pointer was moved across the surface of the participant’s head to collect a minimum of 200 points to calibrate the surface of the participant’s head with the TMS software. The landmark and surface registration steps were verified by comparing the positions of the landmarks and surfaces in real space with the respective locations on the virtual brain scan. A root mean squared error of less than 4.0 mm was considered acceptable for coil placement.

Following the Neuronavigation procedure, a chin support connected to the procedural chair was locked in place to support the participant’s head and minimize head movement throughout the experiment. The TMS coil was placed over the forearm motor hotspot of the participant using the Neuronavigation system. Proper placement of the TMS coil was confirmed by sending three TMS pulses to the motor cortex (intensity: 80%) and observing whether motor evoked potentials were present. If there were no motor evoked potentials, the TMS coil was repositioned until motor evoked potentials were observed. The TMS stimulator (MagPro X100, Magventure) settings were set to biphasic waveform, single pulse mode, and normal current direction for all the TMS pulse stimulations.

Following TMS coil placement, the position and orientation of the TMS coil was set in the Localite software and head movement monitoring was enabled. Throughout the remainder of the experimental session, any movement or head tilt by the participant greater than 6 mm or 6 ^*°*^ triggered an acoustic beep from the software. The participant was instructed to move their head back into place. If they could not find the initial head location, the TMS coil was repositioned by the experimenters. Stimulation marker locations were saved with the position and orientation of the TMS instrument coil each time a TMS pulse was applied. The full experimental setup is displayed in Fig. 3.

**Figure 3.**
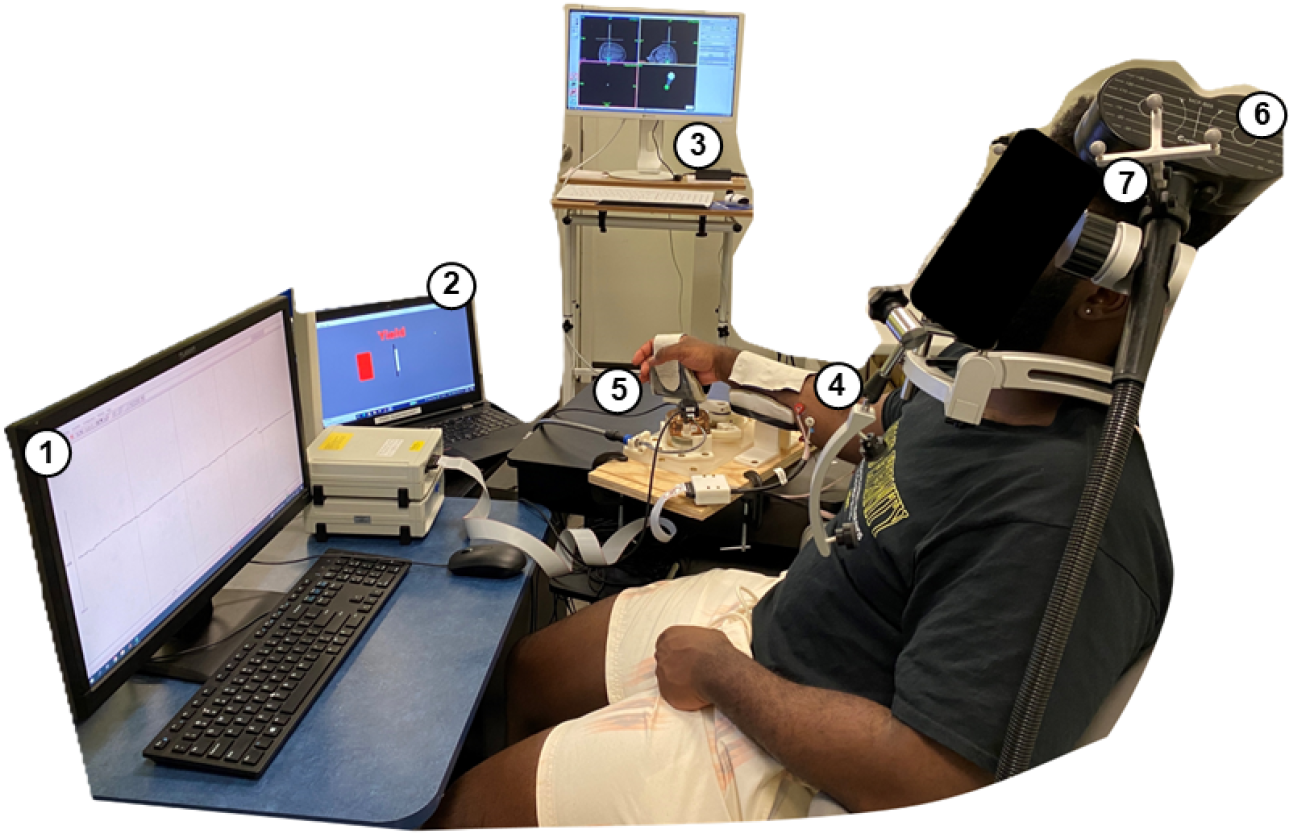
A) Laboratory experimental setup with the TMS system, MR-StretchWrist robot, and FCR surface EMG acquisition. 1-EMG Monitor; 2-2D Visual Display;3-TMS Neuronavigation Computer;4-Surface Electrodes;5-MR-StretchWrist Robot;6-TMS Coil;7-Reflective Markers. The TMS coil is placed on the surface of the head and directed towards the forearm motor hotspot area. Accurate placement of the TMS coil is confirmed with the TMS neuronavigation software, Localite.

## Procedures

### MRI Structural Scan

During a first visit, we acquired a high-resolution structural brain scan for each participant using a Siemens 3T MRI scanner (Center for Biomedical & Brain Imaging, University of Delaware). Participants were screened with an MRI safety screening form to ensure they were safe to be scanned in the MRI. The participant laid in the supine position on the scanner bed and was instructed to relax and minimize movement during the scan. The high resolution T_1_ MP-RAGE structural scan required a total of 30 minutes to perform with a 0.7×0.7×0.7mm^3^ voxel resolution, TR = 2300 ms, TE = 3.2 ms and 160 slices with 256 × 256 px per image.

### Active Motor Threshold and T_MEP_ Determination

Once it was confirmed that the TMS coil was placed over the forearm hotspot of the left motor cortex, we determined the participant specific active motor threshold (AMT) and latency (T_MEP_). The AMT was defined as the minimum TMS intensity (% MSO) that elicited at least five out of ten motor evoked potentials (MEPs) when the muscle was in the target activation state. A MEP was recorded if the measured EMG response exceeded an EMG threshold based on the muscle activation signal measured during isometric wrist contractions. The MEP threshold was estimated on a participant-by-participant basis as the floor raw EMG value of the voluntary contraction signal. This method for determining the active motor threshold has been previously recommended for TMS motor protocols (49).

The participant was cued to maintain a wrist flexion torque of 0.3 N·m as indicated by a visual display, corresponding to roughly 5-10% maximum voluntary contraction (50). The visual display composed of a horizontal bar whose length was mapped to the measured wrist flexion torque, and whose color changed from gray to green if the measured torque was within 0.07 N·m of the target. Once participants matched the target torque, a single TMS stimuli was applied. The TMS intensity started at 40% of the maximum stimulation output (MSO) of the TMS stimulator. If an MEP was observed, the TMS intensity decreased by 10%, otherwise it increased by 10%. The participant was cued to relax following the TMS stimuli at one intensity. Once an MEP was observed, the TMS intensity decreased by 5%. Then, if an MEP was not observed, the TMS intensity was increased by 2% and ten single pulse stimuli were fired with an approximate inter-stimulus time of 5 s. Additional TMS intensity changes were in increments of 2% until we found the lowest TMS intensity that yielded at least 5 MEPs.

The measured value of AMT was used to define stimulation amplitudes in the later portion of the experiment. TMS timing was calculated for the later portions of the experiment based on the participant’s T_MEP_ latency. The T_MEP_ latency was defined as the average delay between the time the TMS was fired and the absolute maximum or minimum of the measured EMG track. T_MEP_ was determined by pooling together the EMG responses elicited by *suprathreshold* stimuli at the participant’s AMT.

### Experimental Task

In the task, the MR-StretchWrist robot was used to elicit stretch reflexes in forearm muscles, TMS was used to stimulate the left motor cortex in proximity of the motor hotspot for muscles in the right forearm, and FCR muscle activity was recorded using surface EMG. Perturbations were applied to the wrist joint in the extension direction to elicit a stretch response in the FCR muscle.

To quantify the effect of single pulse TMS intensity on the TMS-induced LLR inhibition and compare with the effect measured in absence of an imposed stretch, we performed an experiment with N=12 participants. The TMS intensities selected were 90% AMT (*subthreshold*) and 130% AMT (*suprathreshold*). In our study, we applied an MEP latency T_*L*_ = 50 ms (see Fig. 4). This latency was chosen such that the ensuing silent window, supposed to be of approximately 150 ms of duration (32, 35), would completely overlap with the LLR portion of the stretch-evoked response. We applied a perturbation (Pert) condition with a perturbation applied without TMS; Back condition with isometric voluntary contraction without a perturbation; TMS condition with TMS applied during isometric voluntary contraction; and a Pert&TMS condition with TMS applied prior to a perturbation. For each condition with TMS, we applied both the *subthreshold* and *suprathreshold* TMS intensity for a total of six experimental conditions. Each mode was repeated 20 times in a randomized order as an event-related design for a total of 120 trials. All TMS pulses were applied with the coil handle pointed posteriorly at a coil orientation of 45^*°*^ relative to the mid-sagittal plane, oriented in the posterior-anterior direction (P-A). This TMS coil orientation has been shown to be the most effective in eliciting motor evoked responses at lower TMS intensities compared to other orientations (51), and can thus be considered the gold standard for excitatory TMS paradigms.

**Figure 4.**
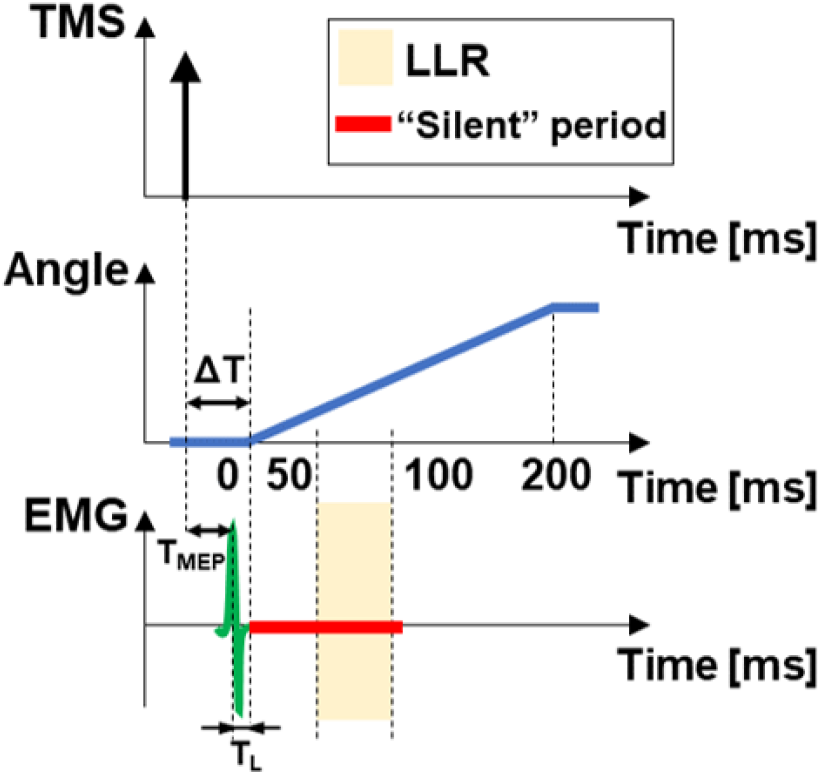
TMS timing diagram showing the synchronization of TMS pulse application, wrist perturbation, expected flexor carpi radialis muscle activity response, and silent period. T_*L*_ is the timing of the peak of the MEP relative to perturbation onset for a *suprathreshold* MEP response condition. The expected muscle activity response is in the case of a *suprathreshold* motor evoked potential.

Prior to the experiment, participants were instructed to ‘Do Not Intervene’ to the perturbation (when applied) as soon as they felt the robot began to perturb them. Each trial began by cueing the participant to maintain a wrist flexion torque of 0.3 N *·* m as indicated by a visual display, just like in the AMT session. Upon matching the torque target, the TMS stimulus would be applied at a delay Δ*R* calculated from a uniform distribution 𝒰_[*a,b*]_ with *a* = 400 ms and *b* = 800 ms. Following perturbation, the robot would return the wrist to the neutral position at a low velocity, and the target bar for the next trial would appear after a delay Δ*T* calculated from a uniform distribution 𝒰_[*a,b*]_ with *a* = 3 s and *b* = 7 s.

## Data Analysis

### EMG

Raw EMG data was acquired for the FCR muscle for all experimental trials and digitized using the EMG amplifier. The digitized EMG data was filtered with a 60 Hz notch filter. The EMG data was demeaned by removing the average EMG signal over time. Then, the EMG signal was filtered by applying a fourth order dual-pass, bandpass filter with a high pass frequency of 20 Hz and a low pass frequency of 500 Hz (9). Then, the EMG response was subject to full-wave rectification by taking the absolute value of the EMG.

Background EMG activity was defined as fully processed EMG signal measured from 125 ms to 75 ms prior to perturbation onset or target disappearing. For a given trial, the processed EMG signal from 200 ms prior to perturbation onset to 250 ms post perturbation onset was subject to statistical analysis. The processed EMG signal was normalized for each trial by dividing it by the average background activity of all trials. Thus, the normalized EMG response in normalized units (n.u.) were obtained for each trial. For each trial, the normalized EMG response from 50 ms to 100 ms post perturbation onset was averaged to obtain the LLR amplitude (LLRa).

### EMG Data Classification by Motor Evoked Potential

Due to the focus of this study on quantifying the effect of TMS intensity on LLR inhibition, EMG signal measured in all TMS ‘ON’ conditions was screened to establish whether a motor evoked potential (MEP) was indeed evoked. To do so, trials were labeled as MEP^(+)^ if the peak of the EMG signal during the expected motor evoked potential window was greater than the mean plus one standard deviation of the distribution of peak rectified background EMG signal measured prior to a perturbation. Trials where the EMG peak was below the threshold were considered MEP^(−)^. Trial classification was performed on an individual participant basis. This EMG data classification method was used for statistical analysis of all experiments. Thus, MEP^(+)^ trials were excluded in *subthreshold* TMS conditions and MEP^(−)^ trials were excluded from *suprathreshold* TMS conditions.

### TMS Target Accuracy Tracking

To confirm validity of our experimental methods, we extracted the TMS marker positions and orientations for each TMS pulse. For each TMS pulse, we obtained the TMS marker location in participant space and the orientation of the TMS coil axes relative to the orientation of the participant space. Using this information, we computed the normal distance from the TMS marker location to the target marker location. We quantified the mean normal distance and standard deviation from the target location to quantify how much each participant had moved during the session as reported in Table 3. To represent the TMS marker location relative to the target location in a compact 2d representation, we computed a plane of reference that contained the target location (origin) and was normal to the average TMS marker location across all TMS stimuli. From this, we plotted the TMS marker locations relative to the target location on a 2D plane. This methodology provides a visualization of the accuracy and precision of TMS stimuli relative to the desired motor hotspot target location.

### Statistical Analysis

#### Participant Specific Stimulation Parameters

Because the threshold for defining an AMT can be inconsistent between studies, we computed the across participant averages and standard deviation for the active motor threshold intensities and T_MEP_, allowing for comparisons with other studies using the same hardware. An expected range for the AMT found by previous studies is between 30% to 50% maximum stimulator output (52) for forearm muscles. The typical MEP latency for hand and wrist muscles has been previously reported to be between 15 ms to 23 ms (53).

**Table 1:** The number of MEP^(−)^ and MEP^(+)^ trials for the *subthreshold* conditions for each participant. Total number of *subthreshold* trials is 40. The count is based on a classification threshold of the mean plus the standard deviation of the peak EMG signal during background contraction.

| Participant ID | MEP (-) | MEP (+) |
| --- | --- | --- |
| P01 | 2 | 38 |
| P02 | 15 | 25 |
| P03 | 19 | 21 |
| P04 | 27 | 13 |
| P05 | 13 | 27 |
| P06 | 19 | 21 |
| P07 | 13 | 27 |
| P08 | 19 | 21 |
| P09 | 15 | 25 |
| P010 | 12 | 28 |
| P011 | 22 | 18 |
| P012 | 24 | 16 |

#### EMG Statistical Analysis

We used a linear mixed model to determine the effects of TMS mode and perturbation on the LLR amplitude. In the model, the fixed effects were Pert mode (2 levels: 0 and 1 for perturbation off and on, respectively), TMS mode (3 levels: No TMS (TMS^(−)^), 90% AMT (TMS^(*sub*)^), and 130 % AMT (TMS^(*supra*)^) and the interaction between Pert mode and TMS mode. The dependent variable was LLRa. No outliers were excluded from the analysis. The linear mixed model included random effects of participant, participant *·* Pert mode, and participant *·* TMS mode.

For the primary statistical analysis, MEP^(+)^ trials were excluded for the *subthreshold* TMS conditions. The statistical analysis was repeated by including all data collected in the *subthreshold* condition, to evaluate sensitivity of the results with respect of mismatch between intended and measured MEP conditions.

In both cases, the mixed model accounted for unequal variance of TMS mode between participants. In presence of a significant fixed effect, post-hoc analysis was performed using Dunnett’s test for multiple comparisons with control to determine whether LLRa in the TMS^*sub*^ and TMS ^*supra*^ modes were significantly different from the control condition (TMS^−^) for both Pert:0 and Pert:1. Beyond comparing LLRa values to baseline, post-hoc tests were conducted to compare the inhibition EMG induced by TMS in presence and absence of an applied muscle stretch, allowing to conclude whether the observed inhibition was specific to a stretch-evoked LLR, or was observed just as a result of an aspecific reduction of background activity. To this aim, we performed the contrasts (TMS^(−)^ − TMS^(*sub*)^)Pert_1_ − (TMS^(−)^ − TMS^(*sub*)^)Pert_0_ and (TMS^(−)^ − TMS^(*supra*)^)Pert_1_ − (TMS^(−)^ − TMS^(*supra*)^)Pert_0_. For all statistical tests, an *α* threshold value of 0.05 was used to determine statistical significance.

## RESULTS

### Participant Specific Stimulation Parameters

The distribution of participant-specific stimulation parameters are reported in Fig. 5. The average AMT was 46.1 %MSO *±* 4.8 %MSO. The average T_MEP_ latency was 22.08 ms *±* 2.3 ms.

**Figure 5.**
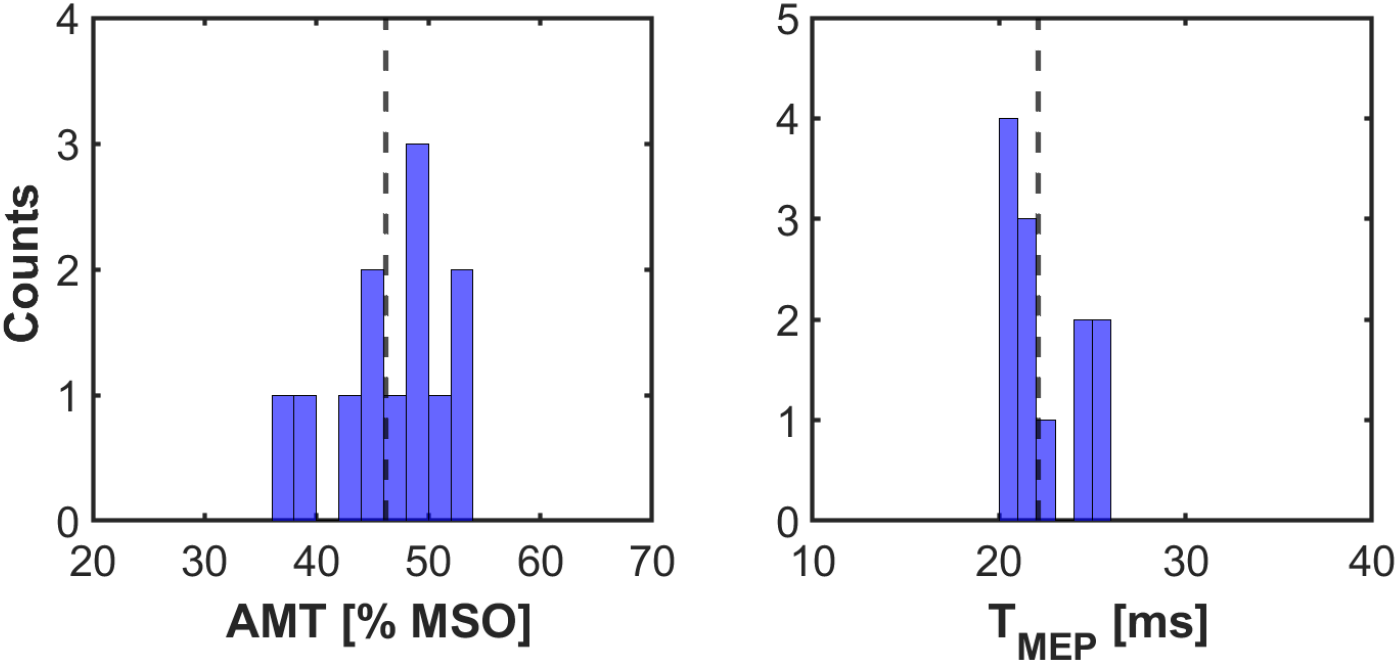
Distribution of active motor threshold and T_MEP_ latency measurements for each group. The dashed black line indicates the mean across participants.

### EMG Timeseries

### Group-level analysis

Experimental EMG time series data and participant specific LLRa are reported in Fig. 6. Data from all trials are plotted in Fig. 6A. All trials that were MEP^(−)^ are plotted in Fig. 6B. Following MEP^(+)^ exclusion, 1 participant had low to no MEP^(−)^ trials and their data were removed from Fig. 6B. It is important to note that no *suprathreshold* trials resulted in MEP^(−)^.

**Figure 6.**
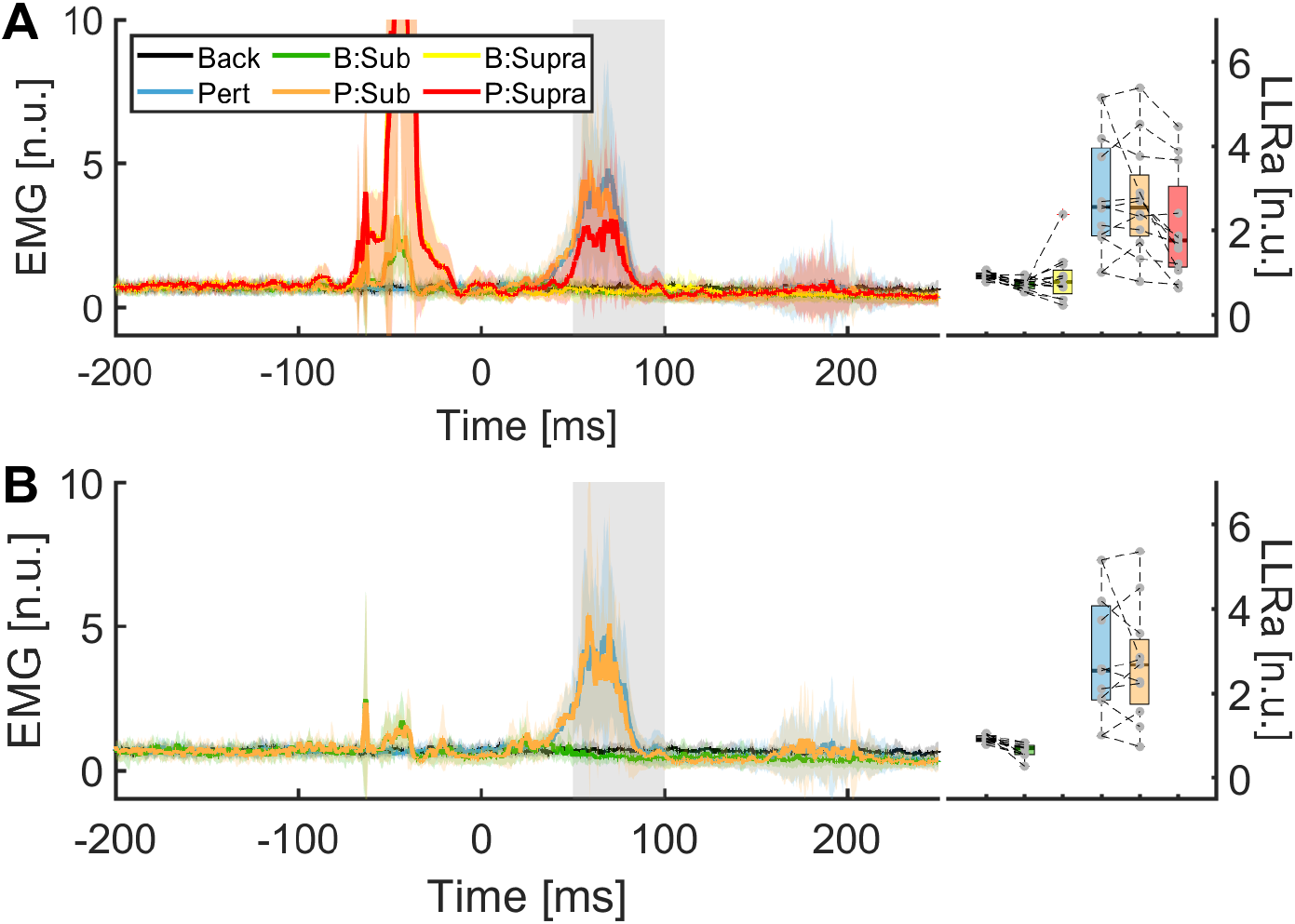
(Left) Timeseries of the normalized EMG signal measured in the different experimental conditions. Group mean normalized EMG responses for each experimental condition are displayed from -200 ms to 250 ms relative to perturbation onset, and a shaded area extends by ± one standard deviation for that condition. (Right) The LLR amplitudes for each experimental condition are displayed. Each dot denotes the within participant average for that experimental condition. The middle line is the median of the distribution with the box ends indicating the 25% - 75% IQR. Whiskers extend out to the furthest datapoint that is not considered an outlier. A: Normalized EMG time series data for all trials. B: Normalized EMG time series data including only MEP^(−)^ trials.

The linear mixed model run on the data after exclusion of MEP^(+)^ trials in the *suprathreshold* TMS condition (primary analysis) identified a significant effect of Pert mode and TMS mode on the normalized LLRa (*F* = 32.21, *df* = 11.0, *p*_*adj*_ <0.001; *F* = 5.49, *df* = 23.3, *p*_*adj*_ = 0.011, respectively), as well as their interaction (*F* = 23.67, *df* = 5564.6, *p*_*adj*_ <0.001) (Table 2). Post-hoc analysis indicated that the protocol consistently evoked long latency responses in the study participants, since LLRa in TMS^(−)^, Pert On trials was significantly greater than in TMS^(−)^, Pert Off trials (2.16 *±* 0.34 n.u., *p*_*adj*_ <0.001). Looking at the effects of the TMS conditions, *suprathreshold* TMS reduced the stretch-evoked EMG signal measured during the LLR window compared to the absence of TMS, since LLRa in TMS^(*supra*)^ trials was significantly smaller than in TMS^(−)^ trials for Pert On (TMS^(*supra*)^ − TMS^(−)^: −0.87 *±* 0.160 n.u., *p*_*adj*_ <0.001). Instead *subthreshold* TMS did not induce a significant effect on the stretch-evoked EMG signal measured during the LLR window, since LLRa in TMS^*sub*^, Pert On trials was not different than in TMS^−^, Pert On trials (TMS^(*sub*)^−TMS^(−)^: −0.41±0.17 n.u., *p*_*adj*_ = 0.060). In absence of a perturbation, both TMS conditions induced a small reduction in EMG signal in the LLRa window, but this reduction was not statistically significant. Specifically, for Pert off trials, LLRa measured in TMS^(*sub*)^ and TMS^(*supra*)^ trials were not significantly different than TMS^(−)^ trials (Pert_0_: TMS^(*sub*)^−TMS^(−)^: −0.29 *±* 0.17 n.u., *p*_*adj*_ = 0.295; TMS^(*supra*)^−TMS^(−)^: −0.06 *±* 0.160 n.u., *p*_*adj*_ = 0.986), see Fig.7.

**Figure 7.**
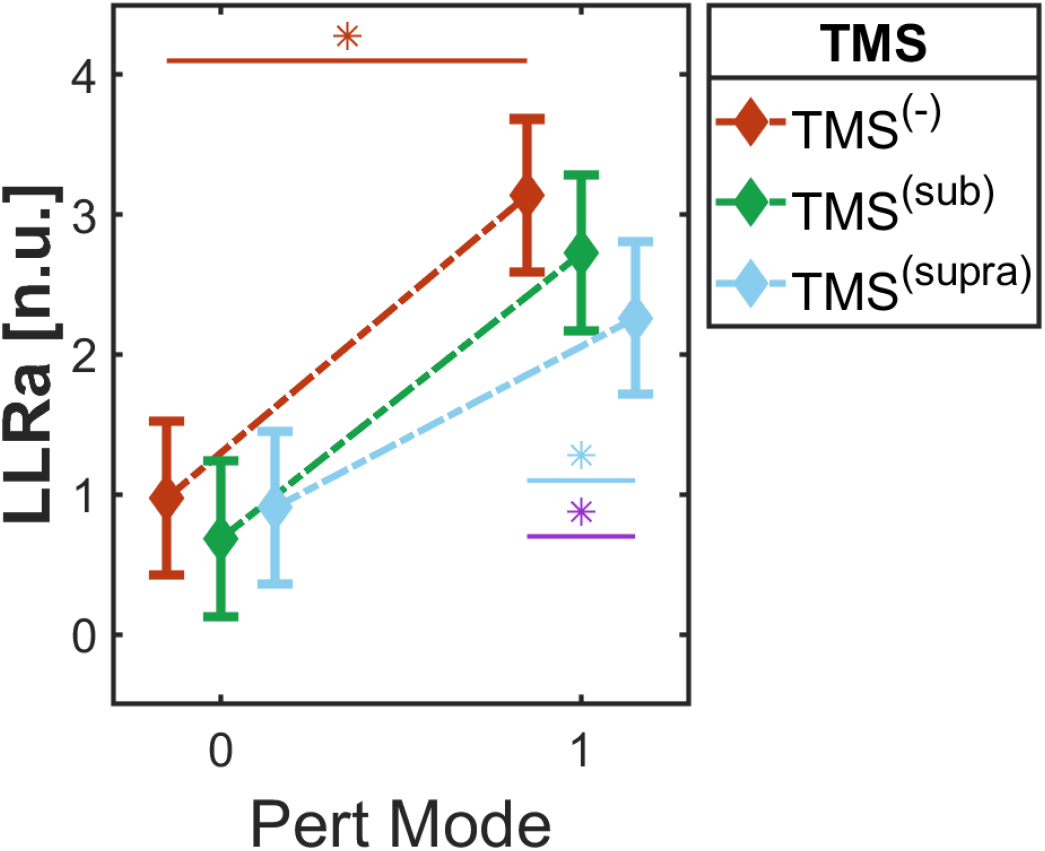
From the experiment task, the effect of different experimental conditions on the LLRa. Asterisks denote significant effects between each experimental condition. Colored asterisk denotes that condition is significantly different compared to the control (Pert:1, TMS^(−)^). Cyan asterisks denotes a significant difference between Pert:1, TMS^(*supra*)^ and Pert:1, TMS^(−)^. Purple asterisks denotes the contrast between ((TMS^(*supra*)^ - TMS^(−)^)Pert_1_ − (TMS^(*supra*)^ − TMS^(−)^)Pert_0_) is significant.

**Table 2:** From the experimental task, the Linear Mixed Model results for the two-way ANOVA test for the fixed effects of Pert Mode (0,1), TMS Mode (TMS^(−)^, TMS^(*sub*)^, TMS^(*supra*)^), and their interaction.

| Fixed Effect | DFNum | DFDen | F Ratio | p-value |
| --- | --- | --- | --- | --- |
| Pert Mode | 1 | 11.0 | 32.21 | <0.001 |
| TMS Mode | 2 | 23.3 | 5.49 | 0.011 |
| Pert Mode·TMS Mode | 2 | 546.7 | 23.67 | <0.001 |

**Table 3:** Average and standard deviation of the normal distance from the TMS marker locations to the target location for each participant.

| Participant ID | Average Distance (mm) | St. Dev. Distance (mm) |
| --- | --- | --- |
| P01 | 2.44 | 0.32 |
| P02 | 5.50 | 0.43 |
| P03 | 7.10 | 0.14 |
| P04 | 6.57 | 0.45 |
| P05 | 5.81 | 0.22 |
| P06 | 5.82 | 0.39 |
| P07 | 4.79 | 2.17 |
| P08 | 2.47 | 0.82 |
| P09 | 1.21 | 0.37 |
| P010 | 0.66 | 0.19 |
| P011 | 3.13 | 0.62 |
| P012 | 3.28 | 0.73 |

The post-hoc contrasts highlighted that an LLR-specific inhibition was observed in the TMS^(*supra*)^ condition, but not in the TMS^(*sub*)^ condition. Specifically, the reduction in LLR in the TMS^(*supra*)^ condition relative to TMS^(−)^ had a significantly greater magnitude in the Pert On condition than in the Pert Off condition ((TMS^(*supra*)^ - TMS^(−)^)Pert_1_ − (TMS^(*supra*)^ − TMS^(−)^)Pert_0_: -0.41 *±*0.063 n.u., *p*_*adj*_ <0.001). Instead, such an LLR-specific effect was smaller and not statistically significant in the TMS^*sub*^ condition ((TMS^(*sub*)^ − TMS^(−)^)Pert_1_ − (TMS^(*sub*)^ − TMS^(−)^)Pert_0_: -0.066 *±*0.083 n.u., *p*_*adj*_ = 0.425), suggesting lack of specificity of the effects with regards to stretch-evoked responses.

The linear mixed model conducted without excluding any data from the TMS^*sub*^ condition provided consistent results with those reported above. The results from this analysis are provided in Table S1 and Fig. S1 in the Supplementary Materials.

### TMS Target Accuracy Tracking

The normal distance from the TMS marker location to the target location was kept within 9 mm throughout the duration of the task for all participants, see Fig. 8. A majority of participants were relatively still and did not move much during the experiment to result in significant mismatches in TMS delivery site. Two participants had larger variability in normal distance during motion due to greater head movement, P07 and P08, nevertheless the normal distance was never greater than 9 mm from the target location. Seven participants stayed at or within 6 mm from the target location during the experimental task, see. Fig. 9. At the group level, the average distance from the TMS marker location to the target location was 4.1 mm *±* 2.1 mm st. dev. On average, participants movement resulted in less than 1 mm displacement from the initial TMS marker location, as quantified by the average within participant standard deviation of the TMS marker locations relative to the target. P010 had excellent TMS coil placement with minimal head movement with less than 3 mm distance for the entire session.

**Figure 8.**
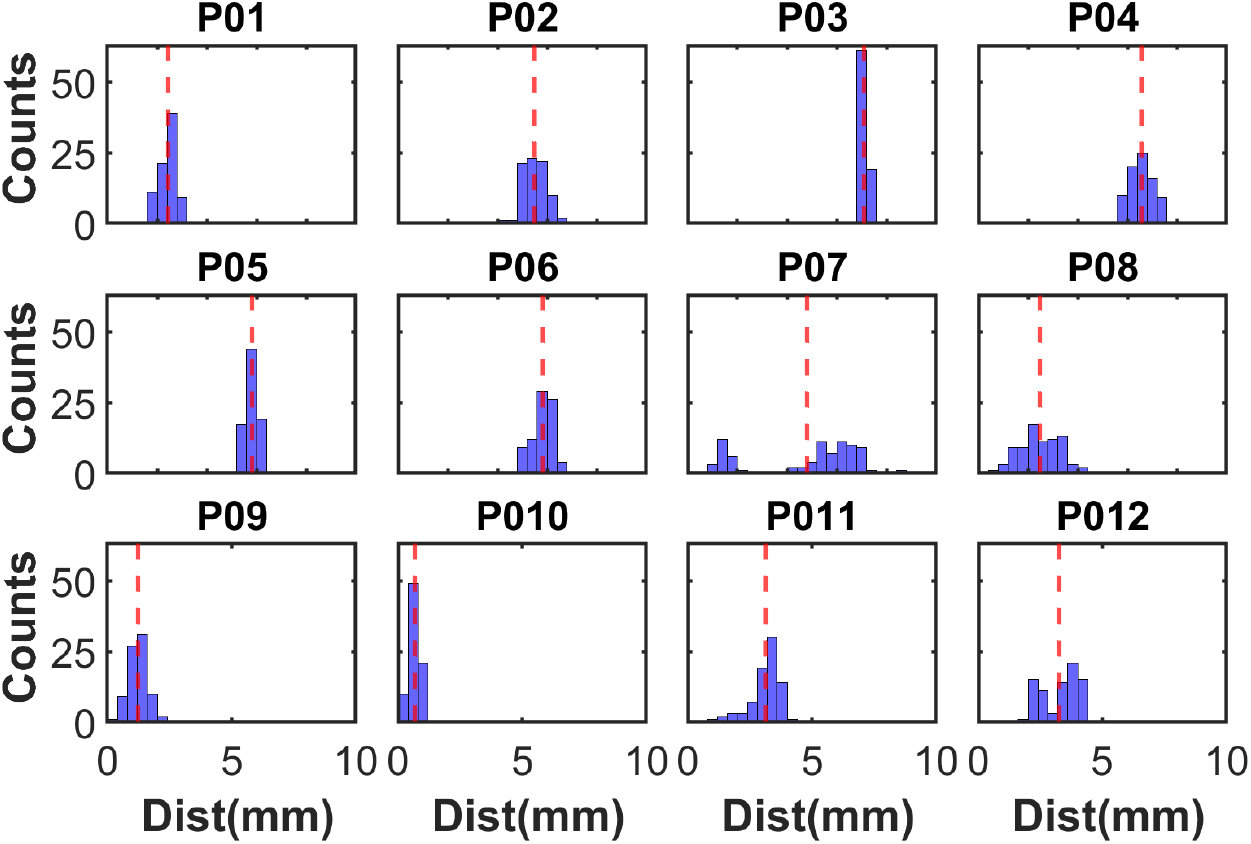
Histogram of the normal distance from the TMS marker location to the target location through the experimental session, broken down for each participant. Red line indicates the mean normal distance to the target for that participant.

**Figure 9.**
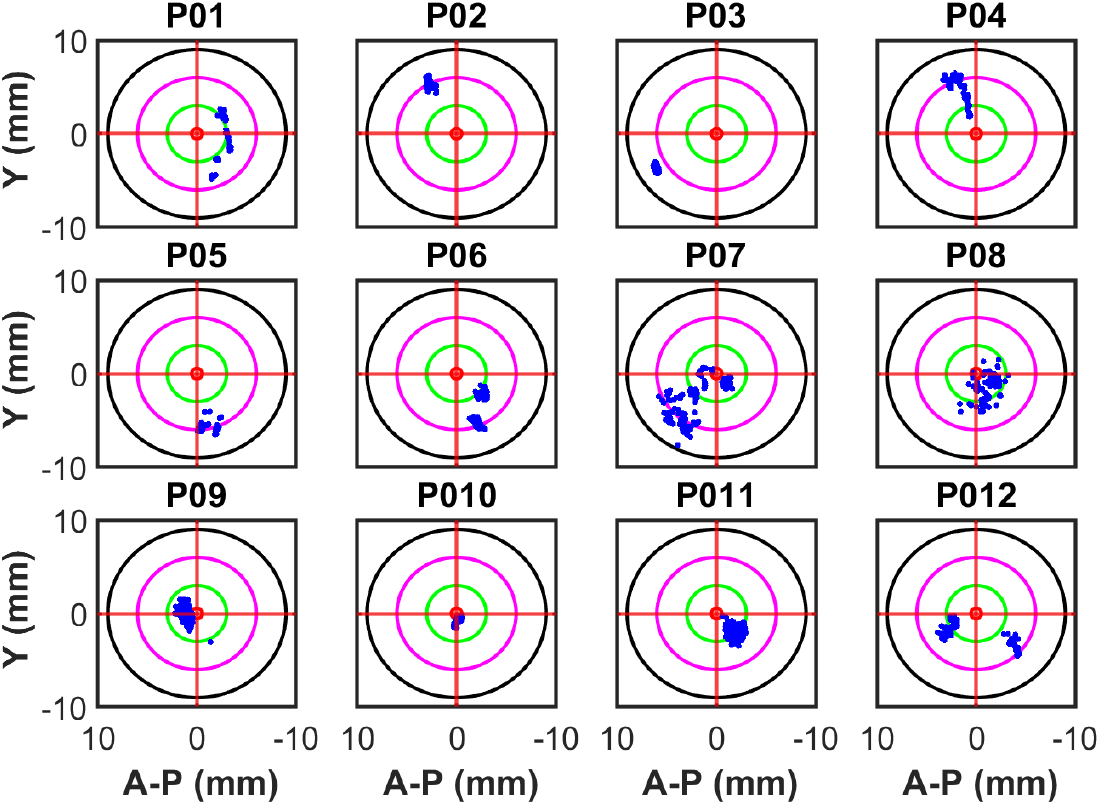
The dispersion of TMS marker locations for each stimulus relative to the target location for each participant. The x-axis is the distance from the target about the posterior-anterior direction. The y-axis is the distance from the target in the medial-lateral direction. Red dot indicates the target location at the origin. Blue dots represent the location of the TMS marker for each TMS pulse applied. The green/magenta/black circles have a radius of 3/6/9 mm, respectively.

## DISCUSSION

The primary objective of this study was to was to determine the effect of TMS intensity on its capability to reduce the CST’s contribution to the LLR in a forearm flexor muscle, FCR. Additionally, we wanted to quantify how accurate our TMS coil setup was in stimulating the forearm motor hotspot of the motor cortex. We used surface EMG to record the LLR in the FCR muscle at different levels of TMS intensity applied prior to perturbation onset. We applied single pulse TMS to the left primary motor cortex to modulate corticospinal excitability during the LLR and voluntary contractions. We used a wrist robot, the MR-StretchWrist, to elicit velocity-controlled perturbations and evoke stretch responses in the FCR muscle.

We found a significant effect of *suprathreshold* TMS intensity on the LLRa compared to the inhibition of voluntary activity during the same time window, suggesting that *suprathreshold* TMS intensities partially inhibit the cortical contribution to a stretch-evoked LLR. We did not find a significant reduction in the LLRa for *subthreshold* TMS pulses. We quantified the normal distance of each TMS pulse to the target location and showed that we accurately and precisely stimulated the forearm motor hotspot with a normal distance of 4.1 mm ± 2.1 mm.

Thus, this study establishes the TMS intensity dependent effect of single pulse TMS when applied prior to perturbation onset on the LLR in the FCR muscle. Knowledge deriving from this study is crucial in understanding how TMS can be used to decouple the contribution of cortical and subcortical components to the long-latency response and provides insight for the design of neuroimaging studies that aim to integrate TMS to quantify brain function associated with LLRs.

### Effect of TMS Intensity on the Long-Latency Response

In this study, we found that *suprathreshold* TMS resulted in LLR specific inhibition, an inhibitory effect that is larger than the one measured in absence of muscle stretch, while *subthreshold* TMS which did not yield any significant inhibition. In presence of a perturbation, we found an overall EMG reduction of 13.1% and 27.8% for the 90% AMT and 130% AMT trials, respectively. While in absence of a perturbation, *subthreshold* TMS resulted in an overall EMG reduction of 29.9% and 7.0% for the 90% AMT and 130% AMT trials, respectively. Generally, we found that a higher TMS intensity resulted in a larger degree of LLR inhibition. This suggests that *suprathreshold* TMS activates the intracortical neurons to specifically inhibit the corticospinal component of the LLR via the cortical silent period. Instead, *subthreshold* TMS did not engage sufficiently such neurons. Of noted, we achieved LLR specific inhibition with 130% AMT, which is a lower TMS intensity than has been previous shown to inhibit the LLR amplitude (163% AMT) when applied prior to perturbation onset. These findings suggest that a motor evoked potential is needed to properly engage and overlay the cortical component of the silent period with the long-latency response.

Our findings align with the degree of inhibition observed in previous studies (43, 44, 4). When applied 50 ms prior to perturbation onset, *suprathreshold* TMS (163% AMT) reduces the LLR amplitude by 38.3% in the biceps brachii and 32% in the ECR for a stiff, do not intervene condition (43, 44). These studies, along with ours, provide empirical evidence for the prolonged cortical silent period that lasts at least 150 ms following the MEP. When TMS is applied during the SLR onset, there is a 15% reduction in the LLR for *subthreshold* TMS and a 43% reduction for *suprathreshold* TMS (4). However, this effect is likely due to the spinal portion of the silent period lasting approximately 50-80 ms post MEP offset.

It is suggested that higher TMS intensities would activate more inhibitory motor neurons and result in a longer silent period duration compared to lower *subthreshold* TMS intensities (54, 55, 56, 57). This may explain why with higher TMS intensities, the silent period duration increases and thus the resulting LLR reduction would be greater over a specified time period.

Prior work has found a significant effect of *suprathreshold* TMS on the LLR modulation in anticipatory or task-dependent modulation. Specifically, Kimura et al., applied TMS prior to a perturbation when the participant had adapted to a force-field following the perturbation. They found that *suprathreshold* TMS applied at the same timing reduces the LLR anticipatory modulation by 60% in the pectoralis major (41).

Spieser et al., applied 120% RMT 50 ms prior to perturbation onset and instructed participants to ‘not react’ or to ‘resist’ the perturbation. They found that *suprathreshold* TMS was capable of significantly reducing the LLR modulation associated with task instruction by 31.5% in the FCR (42). Interestingly, these experiments did not find a reduction of the LLR amplitude itself. Future work could further investigate the TMS intensity dependent effect on task instruction or external forces to determine whether *subthreshold* TMS is capable of reducing LLR modulation.

Though not the main focus of our study, we did find a reduction of EMG activity in all TMS conditions, immediately following the MEP which is due to the initial spinal inhibition. This aligns with the findings from previous work that show the capability of TMS to inhibit the LLR when timed to arrive post-perturbation onset but before SLR onset (4, 39, 40). Given what we know about the timing of the TMS-induced silent period (35), it is reasonable to assume that these paradigms achieved LLR-specific inhibition because they aligned the TMS pulse with the phase where the induced inhibitory effect is spinal, rather than cortical, as sought by our stimulation paradigm.

While we did not find a significant effect of TMS on voluntary activity, we did observe a relative decrease in the voluntary muscle activity, especially in the *subthreshold* trials, during the same time window as the LLR, suggesting that TMS leads to a decrease in voluntary activity. Notably, we noticed a greater reduction in voluntary activity following *subthreshold* than *suprathreshold*. This could be explained by a rebound of EMG activity following the silent period which is greater in *suprathreshold* than *subthreshold* TMS. While participants were instructed to ‘not to intervene’ with the perturbation, it is possible that some participants were startled by the *suprathreshold* TMS pulse and twitched accordingly causing an increase in muscle activity following TMS in some of the background trials.

### Accuracy of TMS Motor Cortex Stimulation

From our TMS marker location tracking, we found TMS pulses consistently stimulated the target area of the motor cortex throughout the experimental session. We found that overall, the normal distance of the TMS marker location from the target location was 4.1 mm on average and deviated from this position by 2.1 mm. As far as we know, this is the first study to report the accuracy and precision of the TMS marker location relative to the target location in the primary motor cortex. This provides a guideline for TMS paradigms to confirm accurate stimulation of the desired brain region and quantify how much individuals move during a TMS experimental session. Head movement tracking ensures that the outcomes we observe are a result of TMS applied to the target location and not an unintended brain region.

### Significance for Future Work

From our results, the LLRa never completely diminishes under any *suprathreshold* TMS latencies, suggesting continued supraspinal contributions during corticospinal inhibition. It has been shown that the LLR has spinal, subcortical, and cortical components (4, 10, 58, 59). As such, the remaining contribution may be due to the sustained contribution of spinal, subcortical, or remaining cortical circuits (4). Future studies need to be done to confirm the effect of corticospinal inhibition on cortical and subcortical structures. Such studies would allow us to investigate whether secondary motor pathways can contribute to functional motor behavior during conditions of reduced corticospinal signal, possibly including those resulting from such lesions responsible for upper motor neuron syndrome. However, the clinical implications of these findings need to be confirmed in a population with corticospinal impairment. Future directions of this work involve performing this TMS protocol with functional neuroimaging to quantify the cortical and subcortical brain areas associated with *suprathreshold* TMS, wrist perturbations, and LLR inhibition.

### Study Limitations

This study has some limitations that should be considered in the discussion of the results. One limitation is that participants may have moved during the experimental procedure relative to the start of the experiment. If the participant moved >6 mm or rotated >6^*°*^, movement may result in stimulating the forearm motor hot spot with a lower TMS intensity than desired resulting in lower cortical inhibition. Alternatively, head movements could result in a better location for stimulating the forearm motor hot spot resulting in a higher TMS intensity than desired. We tracked head movement throughout the experiment and the active motor threshold determination procedure. If the participant moved >6 mm in any direction or rotated >6^*°*^ in any plane relative to the starting position and orientation, the TMS coil was re-localized to be placed over the forearm motor hotspot and the AMT was confirmed. In order to minimize participant head movement, participants were given a chin support to support the weight of their head and keep it in position during the experiment.

Another limitation is the possibility that no stretch responses occurred in the forearm during wrist perturbations or the stretch responses were not large enough compared to background contraction activity. This could be because participants might not have been activating their FCR muscle enough during background contraction or the velocity of the perturbation was not large enough to induce a high stretch response in that muscle. On the other hand, it is possible that some participants could have resisted some perturbations. When resisting a perturbation, there will be a higher long-latency response than when yielding to a perturbation. Ideally, this would not have occurred in all perturbations but may have occurred in some. This would result in the inhibitory effect of TMS not being appropriately quantified in the long-latency response amplitude. To try to minimize the potential for participants to resist perturbations, they were cued to ‘Not to intervene’ to the perturbation before the start of each trial. We found two participants who had low stretch-related motor responses in the Pert:1 level compared to the Pert:0 level. We excluded these participants and re-ran the statistical analysis and found the results hold true even when removing participants with low long-latency responses.

## Conclusions

The objective of this study was to determine was to determine the effect of TMS intensity on its capability to reduce the CST’s contribution to the LLR in a forearm flexor muscle, FCR. We found that *suprathreshold* TMS is capable of inhibiting the cortical contribution to a stretch-evoked LLR, while *subthreshold* TMS did not. Furthermore, during the LLR period following TMS, there remains a positive LLRa suggesting the involvement of remaining cortical, subcortical, or spinal circuitry even in presence of reduced corticospinal tract signal. Further studies need to be done to determine the brain regions associated with the cortical silent period induced by TMS during the LLR. Additionally, future work needs to be conducted to determine the role of subcortical brain regions in the contribution to the LLR during the silent period. Knowledge from these studies could provide further information on how TMS modulates corticospinal excitability and how subcortical brain regions contribute to functional motor output during reduced corticospinal excitability. The findings of these studies would provide crucial information on the role of secondary motor pathways such as the reticulospinal tract on the generation of motor commands during reduced cortical input.

## Acknowledgements

This research was funded by Delaware CTR ACCEL and the Delaware INBRE program.

## Data Accessibility

The data in this work can be made available upon reasonable request.

## Supplementary Materials

### Supplementary Methods

#### Group Level Analysis - All Data

We used a linear mixed model to determine the effects of TMS mode and perturbation on the LLR amplitude. In the model, the fixed effects were Pert mode (2 levels: 0 and 1 for perturbation off and on, respectively), TMS mode (3 levels: No TMS (TMS^(−)^), 90% AMT (TMS^(*sub*)^), and 130 % AMT (TMS^(*supra*)^) and the interaction between Pert mode and TMS mode. The dependent variable was LLRa. No outliers were excluded from the analysis. The linear mixed model included random effects of participant, participant *·* Pert mode, and participant *·* TMS mode. The mixed model accounted for unequal variance of TMS mode between participants.

#### Normalized MEP Peak Calculation

To further investigate the relationship between MEP peak and the LLR inhibition, we computed the normalized MEP peak for each TMS ‘ON’ trial. On a participant-specific basis, we computed the MEP peak as the maximum rectified EMG signal from 0 ms to 50 ms following TMS pulse application. We normalized the MEP peak by dividing the MEP peaks by the average LLRa for each participant. For trials without TMS, we computed the normalized EMG peak signal from -50 ms to 0 ms prior to perturbation onset.

Effect of Normalized MEP peak and LLR amplitude We performed a linear mixed model between the normalized MEP peak and the LLRa to determine whether the normalized MEP peak could significantly explain the variance in the LLRa. The model included fixed effects of Pert(0,1) and Normalized MEP Peak as a continuous variable, as well as their interaction. We accounted for random effects of participantID, participantID*Pert, and participantID*Normalized MEP peak. We computed the indicator parameterization estimates to compare the slope of the Pert:0 to Pert:1. The difference estimate of the slopes and the standard deviation were reported. We further broke down the model by Pert mode to determine whether normalized MEP peak had an effect on the LLRa separately for each pert level. We ran a linear mixed model with fixed effects of normalized MEP peak and random effects of participantID and participantID*normalized MEP peak.

### Supplementary Results

#### Group Level Analysis - All Data

**Table S1:** From the experimental task, the Linear Mixed Model results for the two-way ANOVA test for the fixed effects of Pert Mode (0,1), TMS Mode (TMS^(−)^, TMS^(*sub*)^, TMS^(*supra*)^), and their interaction.

| Fixed Effect | DFNum | DFDen | F Ratio | p-value |
| --- | --- | --- | --- | --- |
| Pert Mode | 1 | 11.0 | 29.76 | <0.001 |
| TMS Mode | 2 | 21.6 | 5.47 | 0.012 |
| Pert Mode·TMS Mode | 2 | 897.1 | 28.99 | <0.001 |

**Figure S1:**
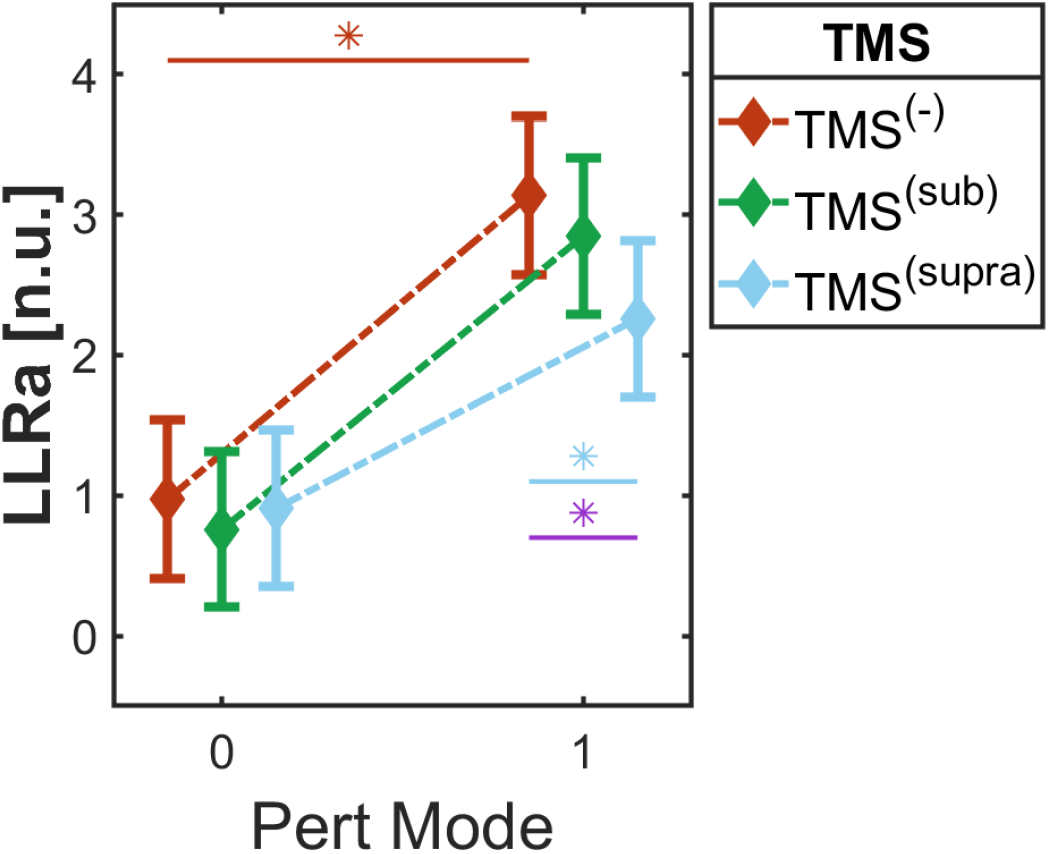
From the experiment task, the effect of different experimental conditions on the LLRa. Asterisks denote significant effects between each experimental condition. Colored asterisk denotes that condition is significantly different compared to the control (Pert:1, TMS^(−)^). Purple asterisks denotes the interaction between Pert:0 and Pert:1 is significant. Cyan asterisks denotes a significant difference between Pert:1, TMS^(*supra*)^ and Pert:1, TMS^(−)^

##### Interaction between Perturbation and MEP Peak

From the MEP peak analysis, we found a significant effect of Pert mode on the LLR amplitude (*p <* 0.001)) (Table. S2). We found a significant interaction between Pert mode and normalized MEP peak (*p* = 0.0356) meaning that the slope of the normalized MEP peak and LLR is significantly different when a perturbation is present compared to when its absent, Fig. S2. The difference estimate between slopes and stan. dev. of the slopes was 0.0005534 *±* 0.000264, meaning that Pert:0 resulted in a less negative slope compared to the Pert:1. When broken down by Pert mode, normalized MEP peak had a significant effect on LLRa for Pert:1 (-0.004 *±* 0.002 n.u., *p* = 0.0386) but not for Pert:0 (0.0004 *±* 0.0006 n.u., *p* = 0.478).

**Table S2:** In the experimental task, the Linear Mixed Model results for the comparison between normalized MEP peak and LLRa for each TMS ‘ON’ mode.

| Fixed Effect | DF Num | DF Den | F Ratio | p-value |
| --- | --- | --- | --- | --- |
| Pert | 1 | 10.8 | 27.57 | <0.001 |
| Normalized MEP Peak | 1 | 5.8 | 1.61 | 0.081 |
| Pert Mode:Normalized MEP Peak | 1 | 915.6 | 5.65 | 0.0356 |

**Figure S2:**
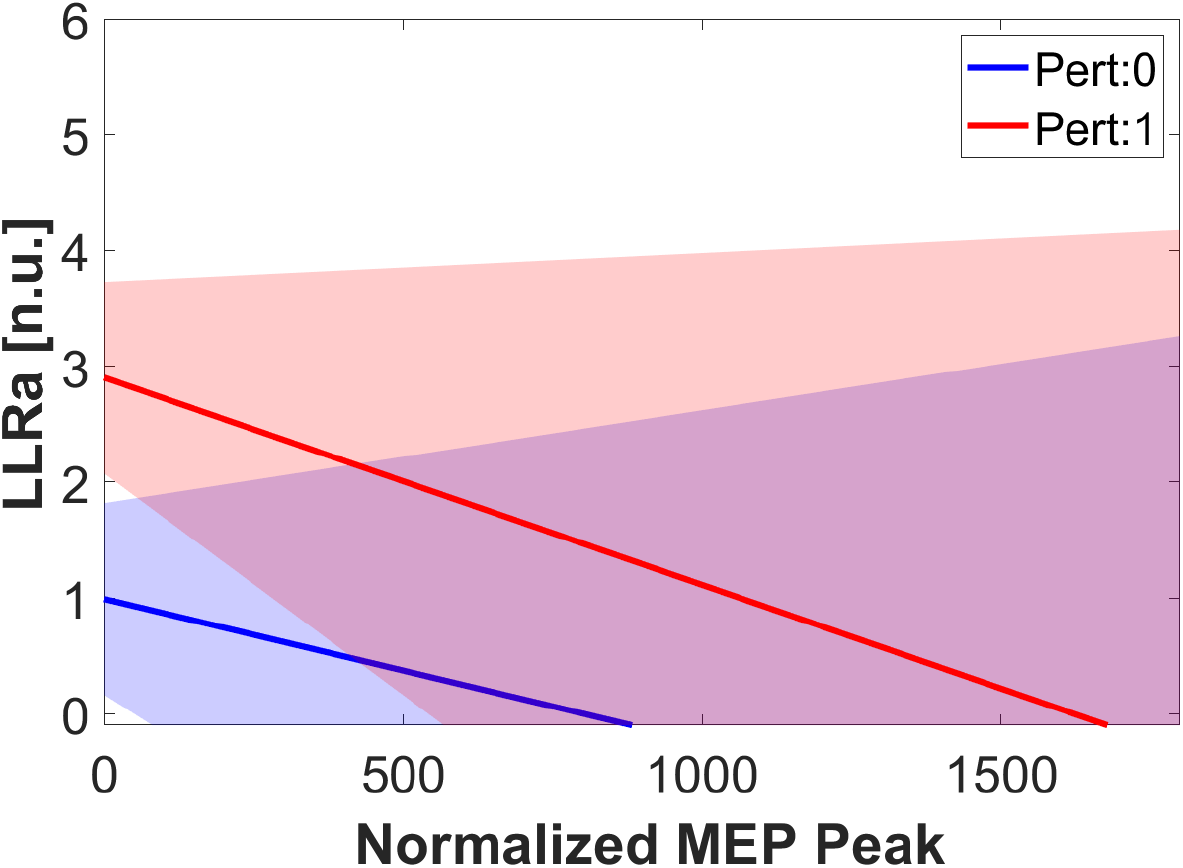
Model estimated relationships and 95% confidence intervals between Normalized MEP peak and Pert Mode on the long latency response amplitude.

#### Relationship between Normalized MEP Peak and Long-latency Response Amplitude

We did not find a significant effect of the normalized MEP peak on the LLRa, meaning that the degree of LLR inhibition did not change with MEP peak. However, we did find a significant effect of the interaction between Pert and MEP peak, meaning that the effect of MEP peak on LLRa depends on whether a perturbation is present. When the perturbation is present, higher MEP peaks result in a greater reduction of the LLRa. It is likely that at higher MEP peaks, the LLR reduction will saturate and at that point, higher MEP peaks will no longer lead to greater LLR-specific inhibition. When there is no perturbation, greater MEP peaks do not result in a greater reduction of the EMG signal. Thus, inhibition of voluntary activity is saturated at lower MEP peaks, while inhibition of stretch-evoked muscle responses continues to increase as MEP peak increases.

